# EXPERIMENTAL DESIGN ISSUES ASSOCIATED WITH CLASSIFICATIONS OF HYPERSPECTRAL IMAGING DATA

**DOI:** 10.1101/2024.08.29.610395

**Authors:** Christian Nansen, Hyoseok Lee, Mohsen B. Mesgaran

**Affiliations:** Department of Entomology and Nematology, University of California, Davis, CA 95616; USDA-ARS, Daniel K. Inouye U.S. Pacific Basin Agricultural Research Center, Hilo, HI 96720; Department of Plant Sciences, University of California, Davis, CA 95616

**Keywords:** Classification models, performance assessment, validation, over-fitting, machine vision

## Abstract

Hyperspectral imaging has emerged as a pivotal tool to classify plant materials (seeds, leaves, and whole plants), pharmaceutical products, food items, and many other objects. This communication addresses two issues, which appear to be over-looked or ignored in >99% of hyperspectral imaging studies: 1) the “small N, large P” problem, when number of spectral bands (explanatory variables, “P”) surpasses number of observations, (“N”) leading to potential model over-fitting, and 2) absence of independent validation data in performance assessments of classification models. Based on simulations of randomly generated data, we illustrate risks associated with these issues. We explore and discuss consequences of over-fitting and risks of misleadingly high accuracy that can result from having a large number of variables relative to observations. We highlight connections of these issues with radiometric repeatability (levels of stochastic noise). A method is proposed wherein a theoretical dataset is generated to mirror the structure of an actual dataset, with the classification of this theoretical dataset serving as a reference. By shedding light on important and common experimental design issues, we aim to enhance methodological rigor and transparency in classifications of hyperspectral imaging data and foster improved and effective applications across various science domains.

## Introduction

Hyperspectral imaging has emerged as a pivotal tool to classify plant materials (seeds, leaves, and whole plants), pharmaceutical products, food items, and many other objects. This communication aims to shed light on two issues in experimental designs that appear to be over-looked or ignored: 1) “Small N, large P” problem (Daee et al., 2016; Mwangi et al., 2014; Zhang and Cao, 2019; Khare et al., 2019), which may lead to model over-fitting (Nansen et al., 2013; Faber and Rajko, 2007; Nansen and Elliott, 2016; Defernez and Kemsley, 1997; Hawkins, 2004). This is a well-recognized phenomenon in machine learning, when the number of spectral bands (explanatory variables, “P”) surpasses the number of observations, (“N”). 2) Absence of independent validation data in performance assessments of classification models. These two issues are relevant across all spatial scales of hyperspectral imaging studies, including benchtop, greenhouse, and airborne applications.

To support our bold claim of these issues being over-looked or ignored, we performed a simple literature search in Google Scholar. Using the search term, “Hyperspectral”, yielded 991,000 results. Adding either the search terms, “Independent validation” (3,910 results), or, ”Model over-fitting” (481 results), shows that >99% of published hyperspectral studies appear not to take these issues into account. As a general rule, model over-fitting is likely to occur when number of explanatory variables (spectral bands in this context), P, is (Defernez and Kemsley, 1997):

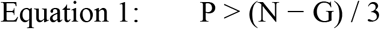

where “N” is the number of observations (pixels or averaged hyperspectral profiles), and G denotes number of treatment classes. As an example, consider a dataset that includes objects divided into three classes, with 12 replicated plants (objects) in each class. In this scenario, the total number of observations, N, would be 36 and concerns about model over-fitting should be raised, if the number of explanatory variables, P, exceeds 11 (equation 1: ((36-3))/3 = 11). Limitation to such a small number of spectral bands presents a significant challenge. If only a few spectral bands are used, then it may be difficult to justify acquisition of hyperspectral data, as potentially unique benefits of hyperspectral imaging may not be fully leveraged. And if a few spectral bands are selected out of an initially large number of spectral bands, then the issue still applies. In some cases, dimensionality reduction methods may be deployed to address the “small N and large P” problem or to eliminate redundant information in highly correlated (collinearity) spectral bands. However, reviews of dimensionality reduction methods highlight that outcomes can be significantly influenced by several factors including (Maaten et al., 2009; Jia et al., 2022; Zebari et al., 2020; Ayesha et al., 2020): 1) potential loss of information, 2) sensitivity of outcomes to the specific methods employed, 3) the computational expense of some methods, particularly those based on optimization, which renders them less feasible for large datasets, 4) the fine-tuning of parameters, which can result in markedly different outcomes, 5) the exacerbation of challenges associated with unbalanced datasets, and 6) the obscured interpretation of hyperspectral imaging data, especially when nonlinear dimensionality reduction methods are used. Thus, it is recommended that dimensionality reduction methods are deployed with caution and accompanied by thorough validation of classification functions.

### Model over-fitting

Most hyperspectral imaging sensors acquire radiometric data in 100-300 spectral bands. Thus, consider the scenario described above but from the opposite direction: estimating the number of observations required if, for instance, a hyperspectral imaging sensor with 200 spectral bands is used in a study involving three treatments. Using Equation 1, we can calculate that 605 whole plants (or other objects) would need to be included in the study, as given by 605 = 3 × 200 + 5. This number of observations far exceeds what is typically feasible under most experimental circumstances, whether in greenhouse plants, petri dishes in growth chambers, field plots, or other contexts. As a result, it is not uncommon to encounter published studies, in which this fundamental guideline or recommendation is either violated or disregarded.

To illustrate the potential consequences of this “small N, large P” challenge, we conducted simulation using random data (see Supplementary Materials for R code). The program generates results from 1,000 simulations based on the following initial parameters:

- Number of observations: Set to 90 by default.
- Number of spectral bands to classify the 90 observations varied and included 1, 10, 20, 30, 40,50, 60, 70, 80, and 90 bands.
- Number of classes: Three, by default. With 90 observations divided into three classes, each class contains 30 observations, ensuring a balanced dataset. This configuration can be adjusted to reflect different proportions matching an actual dataset.
- Reflectance values (mean_val): Assumed to range from 0 to 1 and to follow a normal distribution, with a default mean of 0.5.
- Standard deviation (sd_val) of theoretical reflectance values: Set at two levels, 0.02 and 0.25.
- Number of simulations: Set at 1,000 by default.

As supervised classification, we used the e1071 library to perform both linear and radial kernel Support Vector Machine (SVM) classifications (Dimitriadou et al., 2008). For each kernel, we assessed model performances based on Kappa values from 10 different ratios of P (number of bands) to N (number of observations), keeping the number of observations fixed at 90 (Fig. 1). Two scenarios were performed for each kernel, in which standard deviations of theoretical reflectance values were set at 0.025 (Fig. 1a and b) and 0.25 (Fig. 1c and d), representing what was considered “high” and “low” radiometric repeatability (see below), respectively.

**Figure 1.**
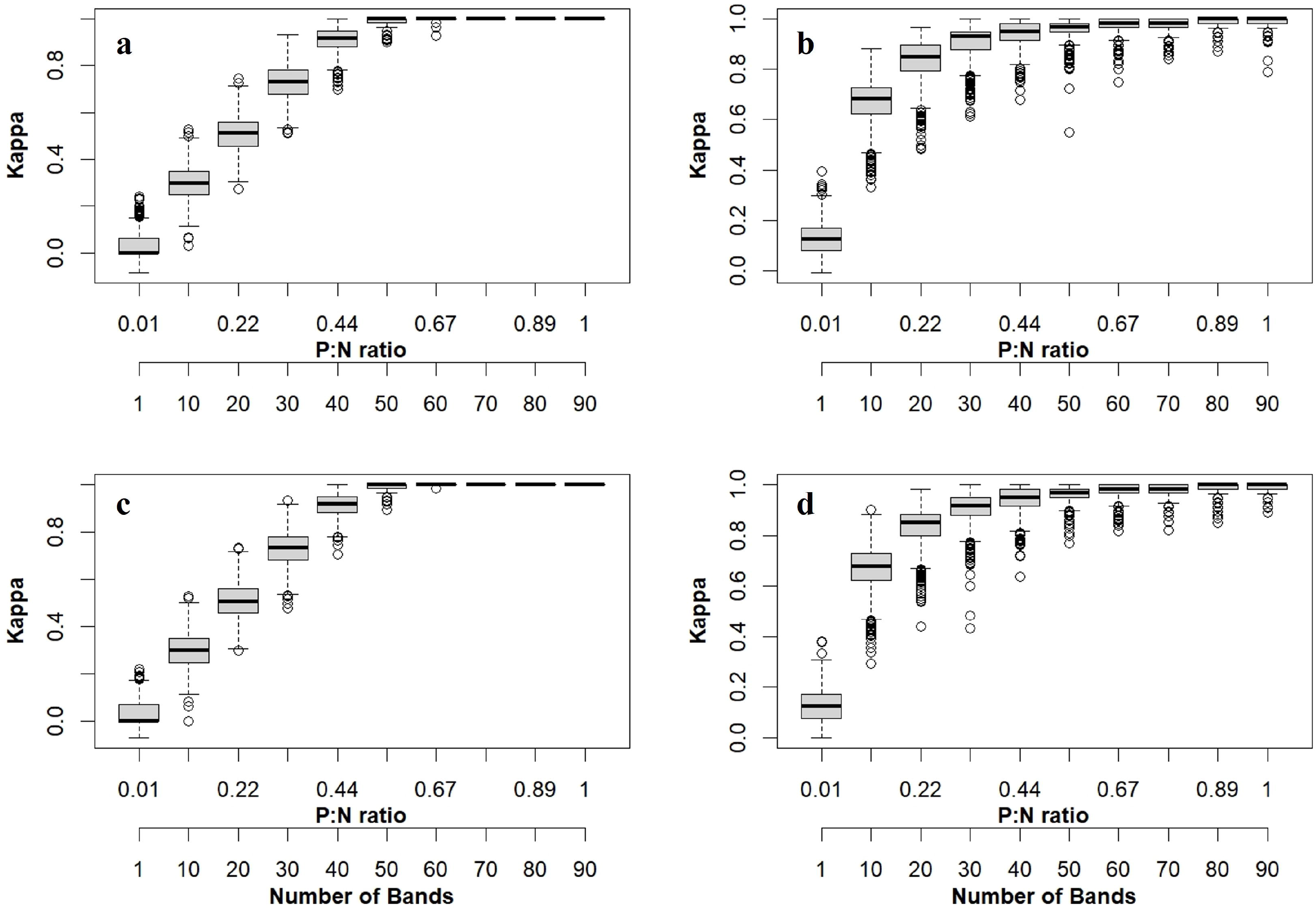
Box plots of Kappa values from 1,000 simulations of random data comprising 90 observations divided into three classes and when classified based on 10 different P:N ratios. Classification results were generated with either high (**a** and **b**) or low (**c** and **d**) radiometric repeatability and with linear (**a** and **c**) or radial (**b** and **d**) kernel functions. Simulated results were generated based on assumption of normal distribution around mean random values of 0.5.

Based linear SVM kernels, similar results were obtained regardless of whether simulated radiometric repeatability was high or low (Fig 1a and c). As general trend, and as expected, classification performance increased progressively with number of spectral bands used as explanatory variables from 1-50. Using only 20 spectral bands yielded Kappa values around 0.5-0.6. Kappa values from classifications with >70 spectral bands were = 1 and did not generate any outliers. Thus with 1,000 simulation, linear kernels generated highly robust and highly accurate classification results. Based on radial SVM kernels, we also observed similar results, regardless of whether simulated radiometric repeatability was high or low (Fig 1b and d). Moreover, average Kappa values exceeded 0.7 when more than 10 spectral bands were used in classification models, regardless of simulated radiometric repeatability level. Compared to linear kernel classifications, radial kernel classifications generated a wider range of classification performances. Meaning, radial kernel functions needed less spectral bands to generate high average classification performance but also appeared to more sensitive than linear kernel functions to unique traits in the random data.

According to Equation 1, the number of spectral bands should not exceed 29 for this dataset (P:N > 0.32) for which our simulations predicted Kappa values between 0.7 - 0.8 and 0.9-1.0 with linear and radial kernel functions, respectively. Regarding interpretation of Kappa values, the following is generally accepted (Landis and Koch, 1977): 0 = poor, 0.01– 0.20 = slight, 0.21–0.40 = fair, 0.41-0.60 = moderate, 0.61–0.80 = substantial, and 0.81–1.00 =almost perfect. In other words, classification performance of 90 observations divided into three categories would be considered “substantial” or almost perfect if classified by random values associated with either low or high spectral repeatability in 30 or more spectral bands.

### Why using simulated data?

A potential critique of using simulated data rather than examining consequences for an actual dataset is two-fold: 1) General framework: We have provided a general framework that can be applied to investigate all datasets. The R code provided as supplementary data can be easily modified to match the structure of any actual dataset, according to: number of observations, number of classes, balance of observations among classes, and number of spectral bands. Furthermore, the proposed SVM classification can be easily replaced by any other classification function. 2) Random data as reference: By generating a random dataset with a structure identical to any actual dataset, classification outcomes can be used as reference in classifications of actual datasets. For example, if classification simulations are performed with random data and result in Kappa values ranging from 0.60-0.80, and classification of an actual dataset yields a Kappa value of 0.80, it suggests that most of spectral traits used for classification in the actual dataset is likely associated with random noise. In essence, use of random data in this exercise is not a limitation but a deliberate methodological choice. Quantifying classification performance of a random data set can greatly inform interpretations of results from actual datasets within the context of recognizing that model over-fitting due to a high numbers of spectral bands is a risk that must be addressed.

### Model validation

We wish to highlight the importance of including independent validation data in performance assessments of classifications. It is surprising to us, and also concerning, that only 3,910 references out of 991,000 scientific references to “Hyperspectral” in Google Scholar included “Independent validation”. This concern is directly connected to a broader issue related to radiometric repeatability of hyperspectral imaging data (Nansen et al., 2023; Vilaseca et al., 2014; Anderson and Peleg, 2007; Peleg et al., 2005). Absence of independent validation data can undermine the reliability of classification models, as it is paramount to assess how models perform when applied to new and independent datasets. In the context of hyperspectral imaging, where data consistency and repeatability are critical, this lack of validation becomes even more significant. Additionally, it is common for authors to collect hyperspectral imaging data at a single time point and/or from one location. Peleg et al. (2005) published an article, that made compelling and logically sound assertions about analyses of hyperspectral imaging data: “Most analysts are either unaware of this problem or just ignore it. Hyperspectral image cubes acquired in consecutive flights over the same target should ideally be identical. In practice, two consecutive flights over the same target usually yield significant differences between the image cubes. These differences are due to variations in: target characteristics, solar illumination, atmospheric conditions and errors of the imaging system proper.” They further stated: “Clearly, when the non-repeatability variance is similar in magnitude to the variance of the spectral or spatial information of interest, it would be impossible to use it for classification or quantification prediction modelling.”. Despite the clarity and validity of these points, this article has garnered only modest research attention, with only 16 citations by the time we prepared this article.

Although their focus was on classifications of hyperspectral imaging data acquired with airborne systems, Peleg et al. (2005) demonstrated that similar repeatability issues, albeit less pronounced, exist with benchtop acquisitions of hyperspectral imaging data under highly controlled laboratory conditions.

Hyperspectral imaging datasets are inherently associated with some degree of radiometric inconsistency, leading to reduced radiometric repeatability. The logical consequence of radiometric inconsistency is a concern that classification models generated based on one dataset may not perform adequately when applied to a new or independent dataset. Therefore, there exists a strong connection between levels of radiometric repeatability in actual hyperspectral datasets and the need for validation procedures. And this is also the reason behind addition of stochastic noise and accompanying simulations were included in the exercise described above.

The frequent absence of independent validation may come as a surprise, underscoring a significant gap in the methodological rigor of many studies. This oversight can have significant implications for the validity and generalizability of the findings. We suggest that studies without any form of validation, should not be published, as they do not allow for adequate assessments of concerns related to model over-fitting. Such an omission compromises scientific integrity and reliability of research findings. The most common validation practice is to perform k-fold cross-validation (Fauvel et al., 2015; Jin et al., 2009; Huang et al., 2019). In k-fold cross-validation, the data is divided into ‘k’ equally (or nearly equally) sized folds or chunks. The model is then trained on ‘k-1’ of these folds and tested on the remaining fold. This process is repeated ‘k’ times, with each fold serving as the test set exactly once. Results from ‘k’ tests are averaged to produce a single estimation of model performance. This method ensures that every observation in datasets is used for validation exactly once, providing a comprehensive assessment of the model’s performance across different data subsets (Hastie et al., 2009). The R code provided in the supplementary materials file performs 10-fold validation (cross = 10), and average classification performance of the random dataset with three classes, irrespectively of number of spectral bands included, yields results close to 33% (the expected outcome when classifying random data in three classes). Thus, k-fold validation is a powerful and often a reliable way to assess classification performance. However, a more refined approach, commonly employed in engineering and computer science, but less frequently in biological sciences, involves training and optimization of classification models on specific datasets, followed by validation on new and independent datasets.

### Basic data constraints

As emphasized earlier, it is vital to include as many observations as possible to minimize concerns about model over-fitting and to fully explore the benefits of hyperspectral imaging. However, due to the second issue discussed above, we are also advocating for reservation of separate datasets for independent validation. We fully recognize that this recommendation creates a conundrum, as practical constraints and the need for compromises in experimental design become apparent.

Balancing desire for comprehensive data exploration with necessity for robust validation requires careful consideration and planning. For example, we previously illustrated how use of a hyperspectral imaging sensor with 200 spectral bands would necessitate the inclusion of 605 plants or observations. If an additional validation dataset were required, then one would face the daunting task of conducting an experiment with about 1,000 whole plants, excised leaves, petri dishes, or other objects!

Maintaining such a large number of objects under nearly identical conditions would likely be a challenging endeavor. Implementing blocking and other measures to control variability might be necessary, further adding to the logistical complexities of experiments and of data analyses. Time required to acquire hyperspectral imaging data from large number of objects can introduce additional errors due to possible variability in imaging conditions, such as, lighting, temperature, and humidity. If hyperspectral imaging data are acquired outside with sunlight as light source, then changes in sun angle, cloud cover, and atmospheric composition will add further noise to datasets and need for deployments of radiometric calibration (Agapiou, 2020; Iqbal et al., 2018; Karpouzli and Malthus, 2003; Smith and Milton, 1999; Aasen et al., 2018; Zarzar et al., 2020; Wang and Myint, 2015; Mafanya et al., 2018; Del Pozo et al., 2014; Poncet et al., 2019; Baugh and Groeneveld, 2008). We acknowledge the challenges associated with adhering to recommendations regarding P:N ratio, along with constraints related to costs, availability of labor and equipment, and time. Despite these hurdles, addressing and embracing these experimental design issues, and openly discussing their potential impacts on study results will likely enhance the integrity and validity of the research.

### Final remarks

There is widespread excitement about use of hyperspectral imaging, to tackle natural science challenges, improve commercial operations, and inspire students to pursue careers in science and engage in multi-disciplinary research projects encompassing ecology, machine learning, engineering, and computer programming. It is with shared enthusiasm and commitment to the field that we have penned this communication aiming to shed light on what seems to be frequently over-looked issues. Possible solutions include, but are not limited to:

Spectral Binning. This involves averaging of spectral bands to reduce their number below the number of observations, thereby mitigating the P:N ratio issue.

Increasing number of observations. By including more observations, the P:N ratio can be improved and lead to enhanced robustness of classifications.

Dimension (feature) reduction. Utilizing techniques like principal component analysis, linear discriminant analysis, or other feature selection methods to reduce the dimensionality of the data. This can help in eliminating redundant or irrelevant spectral bands, making classification models more interpretable and less sensitive to over-fitting.

Multiple time points or site acquisition. Acquiring hyperspectral imaging data from the same objects at different time points or locations can provide comprehensive insights in levels of radiometric repeatability.

High spatial resolution data acquisition. Acquisition of high numbers of hyperspectral pixel profiles from a single seed, leaf, or plant. Subsequent random subsampling of hyperspectral pixel profiles from individual objects can be performed. However, pseudo-replication should be a major concern with this approach, and it should be applied with caution and accompanied by thorough validation of classification models (Peleg et al., 2005).

Independent validation. Due to potential concerns about radiometric repeatability (Nansen et al., 2023), a separate and truly independent dataset should be set aside and used for independent validation of classification models.

These strategies, while offering potential solutions, also bring about their own complexities and considerations. The balance between the number of spectral bands, observations, and the need for validation presents a multifaceted challenge that requires careful consideration and planning. By embracing these challenges and thoughtfully addressing their potential effects on study results, the scientific community can foster a more rigorous and transparent approach to hyperspectral imaging.

## Data availability

The datasets generated and/or analysed during the current study are not publicly available due to the fact that only theoretical data are included. However, R code for generation of theoretical data is provided as a supplementary file. Readers can generate their own data, using either default settings in the provided R code, or they can adjust the code so that theoretical data match/align with specific data sets. Alternatively, theoretical data are available from the corresponding author on reasonable request.

