## Supplementary material for "EXPERIMENTAL DESIGN ISSUES ASSOCIATED WITH CLASSIFICATIONS OF HYPERSPECTRAL IMAGING DATA": R code

**Supplementary materials**

\# Load the required library

library(e1071)

\# Initialize parameters

num_observations <- 90

band_numbers <- c(1, 10, 20, 30, 40, 50, 60, 70, 80, 90)

num_classes <- 3

num_simulations <- 1000

mean_val <- 0.5

sd_val <- 0.02

\# Initialize results storage

results_list <- list()

\# Define class probabilities

class_probs <- c(0.33, 0.33, 0.33)

\# Loop through each band number

for (P in band_numbers) {

kappa_values <- numeric(num_simulations)

\# Run simulations

for (i in seq_len(num_simulations)) {

\# Generate random dataset

random_data <- matrix(rnorm(P * num_observations, mean_val, sd_val), nrow = num_observations)

colnames(random_data) <- paste0('B', seq_len(P))

\# Add Class column

class_labels <- paste0('C', sample(seq_len(num_classes), num_observations, replace = TRUE, prob = class_probs))

random_data <- data.frame(random_data, Class = factor(class_labels))

\# Perform SVM classification

svm_result <- svm(Class ~ ., data = random_data, cross = 10, kernel = "linear")

\# Calculate Kappa statistic

pred <- predict(svm_result, newdata = random_data)

confusion_matrix <- table(random_data$Class, pred)

kappa_values[i] <- classAgreement(confusion_matrix)$kappa

}

\# Store Kappa values and P:N ratio

results_df <- data.frame(Kappa = kappa_values, P_N_ratio = round(P / num_observations, 2))

results_list[[as.character(P)]] <- results_df

}

\# Combine results into one data frame

final_results <- do.call(rbind, results_list)

\# Create boxplot

unique_P_N_ratios <- unique(final_results$P_N_ratio)

positions <- seq(1, length(unique_P_N_ratios))

old_par <- par(mar = c(6, 4, 2, 2))

bp <- boxplot(Kappa ~ P_N_ratio, final_results, ylab = '', xlab = '')

mtext("Kappa", side = 2, line = 2.5, font = 2)

mtext("P:N ratio", side = 1, line = 2, font=2)

mtext("Number of Bands", side = 1, line = +5, font=2)

axis(side = 1, at = positions, labels = band_numbers, tick = TRUE, line = +3)

par(old_par)
